# Negative affect homogenizes and positive affect diversifies social memory consolidation across people

**DOI:** 10.1101/2023.02.20.528994

**Authors:** Siddhant Iyer, Eleanor Collier, Emily S. Finn, Meghan L. Meyer

**Affiliations:** Columbia University, New York, NY, 10027; University of California, Riverside, CA, 92521; Dartmouth College, Hanover, NH, 03755

## Abstract

We are often surprised when an interaction we remember positively is recalled by a peer negatively. What colors social memories with positive versus negative hues? We show that when resting after a social experience, individuals showing similar default network responding subsequently remember more negative information, while individuals showing idiosyncratic default network responding remember more positive information. Results were specific to rest after the social experience (as opposed to before or during the social experience, or rest after a nonsocial experience). The results provide novel neural evidence in support of the “broaden and build” theory of positive emotion, which posits that while negative affect confines, positive affect broadens idiosyncrasy in cognitive processing. For the first time, we identified post-encoding rest as a key moment and the default network as a key brain system in which negative affect homogenizes, whereas positive affect diversifies social memories.

## Introduction

Multiple people often partake in the same social event. Yet people may end up with different memories of the shared experience. Friends are often surprised to learn that a conversation one of them remembers going well is recalled by their friend as unsuccessful. Moreover, interpersonal memories are rarely “neutral” and instead intertwined with affect. When we recall our times spent with others, we often recall our interaction partners’ affect, how they made us feel, and/or the overall sentiment of the experience. Yet memory research has not fully determined how social experiences translate into affect-laden interpersonal memories that vary across people. Filling this gap is not only critical to the basic science understanding of real-world learning and memory. It also has implications for understanding mental health, given that affective memory biases are implicated in multiple psychiatric conditions.^1,2,3^

Consolidation mechanisms at rest by the brain’s default network may explain the individualized affect stamped onto interpersonal memories. Extensive research suggests memories are consolidated (i.e., codified in memory) during rest after an experience^4^ (i.e., post-encoding rest). Default network functional connectivity (i.e., timecourse correlations of neural activity between regions) during post-encoding rest, in particular, predicts more accurate memory for people, such as the association between a person’s face and their occupation^5^. Further, while affective reactions (e.g., high arousal) are linked to limbic regions outside of the default network, the mental representation of affective reactions (e.g., the construal of a high arousal state as tense versus excited) is associated with default network regions.^6,7,8^ Moreover, individuals with similar beliefs (e.g., political partisans) demonstrate neural synchrony in default network regions while processing belief-relevant stimuli,^9,10,11,12,13^ further hinting at the possibility that default network consolidation processes may help explain how some people end up with similar (vs. dissimilar) affective content in their memories.

To investigate this possibility, we had subjects watch videos of people discussing a sensitive topic: their experience with cystic fibrosis, in which they shared negative information (e.g., their challenging symptoms) as well as positive information (e.g., how their experience has helped them value their meaningful relationships). Using videos of cystic fibrosis patients allowed us to test our hypotheses in a context where individual differences in positive versus negative affect meaningfully predict well-being. That is, it has been shown that focusing on the positive meaning in a patients’ experience promotes mental health in both patients and caretakers.^14,15^ As a control condition for the interpersonal content shared in the patient videos, participants also watched videos describing the biology of cystic fibrosis (e.g., the genetic basis of cystic fibrosis). Before and after watching the videos, participants completed resting state scans and subsequently completed a surprise memory test, in which they wrote down everything they remembered from the videos. Participants’ memories were submitted to a sentiment analysis that quantified negative and positive content. We were therefore able to test whether individual differences in default network consolidation predicted individual differences in the affect incorporated into social memories.

Importantly, past research suggests negative and positive affect color cognition asymmetrically, such that positive evaluations are often more idiosyncratic than negative evaluations.^16,17,18^ In fact, the “broaden-and-build” theory of positive emotion suggests that while negative affect constricts the nuance of cognitive processing (i.e., generates prototypical thought patterns), positive affect diversifies the types of thoughts entertained^19,20^ (i.e., generates idiosyncratic thought patterns). While the broaden-and-build theory has not been directly tested with neural measures, one prior study provides partial support; it found negative affect increases similarity in default network neural responding across participants during video watching.^55^ Yet, whether positive affect diversifies neural responses remains untested, as does the possibility that broaden-and-build effects occur during post-encoding rest to meaningfully predict individual differences in memory.

The broaden-and-build perspective, in conjunction with the consolidation literature, predicts that people with a negative bias should engage in similar, whereas those with a positive bias should engage in unique, default network consolidation processes. To test this possibility, we used an “Anna Karenina” model^21^, so named after the opening line of Tolstoy’s famous novel, which goes, “All happy families are alike; each unhappy family is unhappy in its own way.” Although Tolstoy’s line posits greater similarity for positive vs. negative experiences, this need not be the case; the Anna Karenina model simply tests whether there is greater similarity in one set of responses compared to another. For the present study’s hypotheses, our Anna Karenina model predicted that subjects with negative memories would exhibit similar neural responses, while those with positive memories would exhibit idiosyncratic neural responses. To examine the specificity of results to consolidation at rest, the Anna Karenina model was tested on each phase of the experiment (baseline rest, video watching, and post video rest). To examine the relative specificity of results to the default network, analyses were repeated separately on limbic regions, as well as across the whole brain.

## Results

### Capturing Individual Differences in Memory Affect

To assess whether affect becomes consolidated into social memories during post-encoding rest, participants completed functional magnetic resonance imaging (fMRI) while undergoing the following experimental phases. Participants watched four videos in which people with cystic fibrosis discussed their negative and positive experiences. This “social encoding” phase consisted of videos from four cystic fibrosis patients, with each patient video lasting approximately 4-minutes. Participants also watched four, approximately 4-minute Khan Academy videos describing the biology of cystic fibrosis (“nonsocial encoding” phase). The order of social and nonsocial encoding was counterbalanced between participants. In between encoding phases, participants completed 6-minute resting state scans: a *baseline* rest scan, *as well as resting state scans after encoding, here termed “social consolidation” and “nonsocial consolidation*.*”* After their scan session, participants completed a surprise free recall test on a computer, in which they viewed a snapshot of each video and were asked to type everything they could remember from the video. The paradigm is depicted in Fig. 1.

**Fig. 1.**
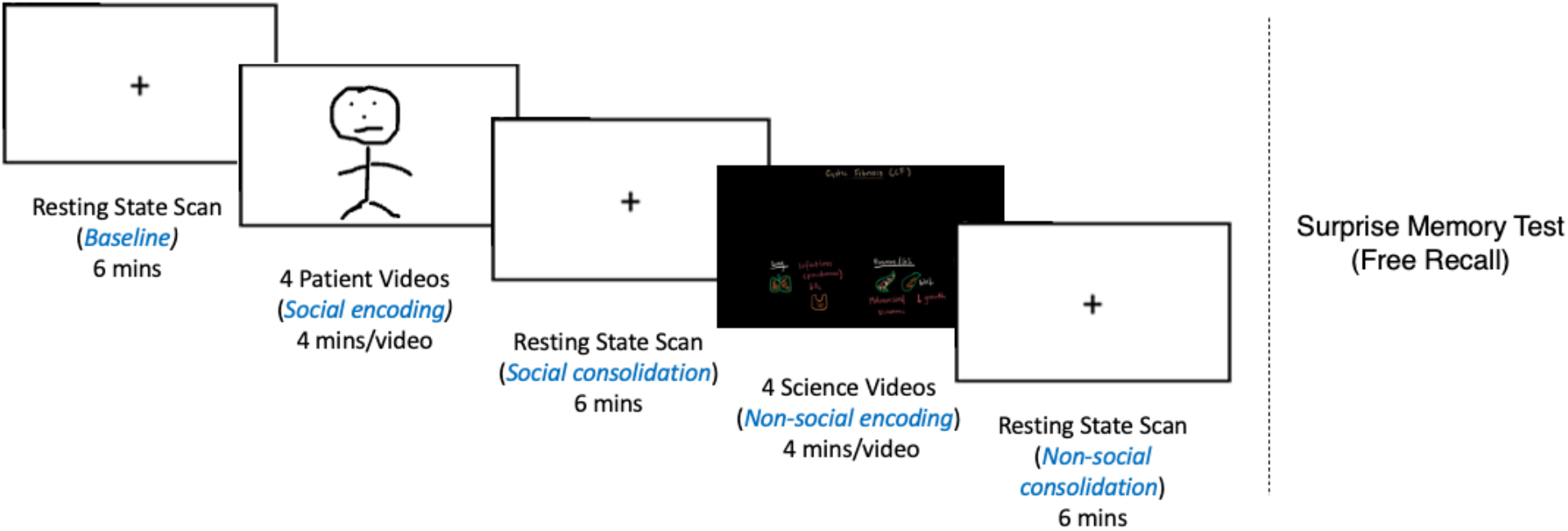
Paradigm. Social and nonsocial encoding were interleaved with resting state scans (i.e., “baseline” and “consolidation” phases). Whether participants first completed social or nonsocial encoding was counterbalanced between participants. After fMRI scanning, participants completed a surprise free recall memory test for which they wrote down everything they remembered from each video. Note that to exclude any identifying information in the manuscript, we have replaced patients’ actual faces with a drawing.

To derive an objective measure of the affective content in subjects’ memories, their written responses to the surprise free recall test were submitted to a sentiment analysis, which is a natural language processing approach used to determine the degree to which language is positive, negative, or neutral. Specifically, we used affect Valence Aware Dictionary and sEntiment Reasoning (VADER^22^)’s Sentiment Intensity Analyzer module, which was created by leveraging and improving upon lexicons and techniques from existing natural language processing models (e.g., LIWC, ANEW), independent Amazon Mechanical Turk word ratings, and machine learning text classifiers.

Fig. 2 depicts the words used by two participants to describe the social videos, one of which had a highly negative affect score (Fig. 2A) and the other a highly positive affect score (Fig. 2B). Negative words are shown in red, positive words are shown in blue; bigger words have stronger affect (highly negative or highly positive). It is noteworthy that participants’ affect scores were not significantly correlated with the number of affective words (*r* = .096, *p* = .554), indicating that affective valence was not meaningfully related to the overall amount of affect expressed in their recall. Further, participants’ affect scores were not significantly correlated with the length of their recall (*r* = .243, *p* = .131), suggesting affect was not confounded with the amount of information recalled. In keeping with the main analyses, all the above tests were computed using spearman correlation, such that inferences could be drawn regarding the *rank order* of subject pairs without assuming linearity between the pairs’ affect scores and connectivity similarity.

**Fig. 2.**
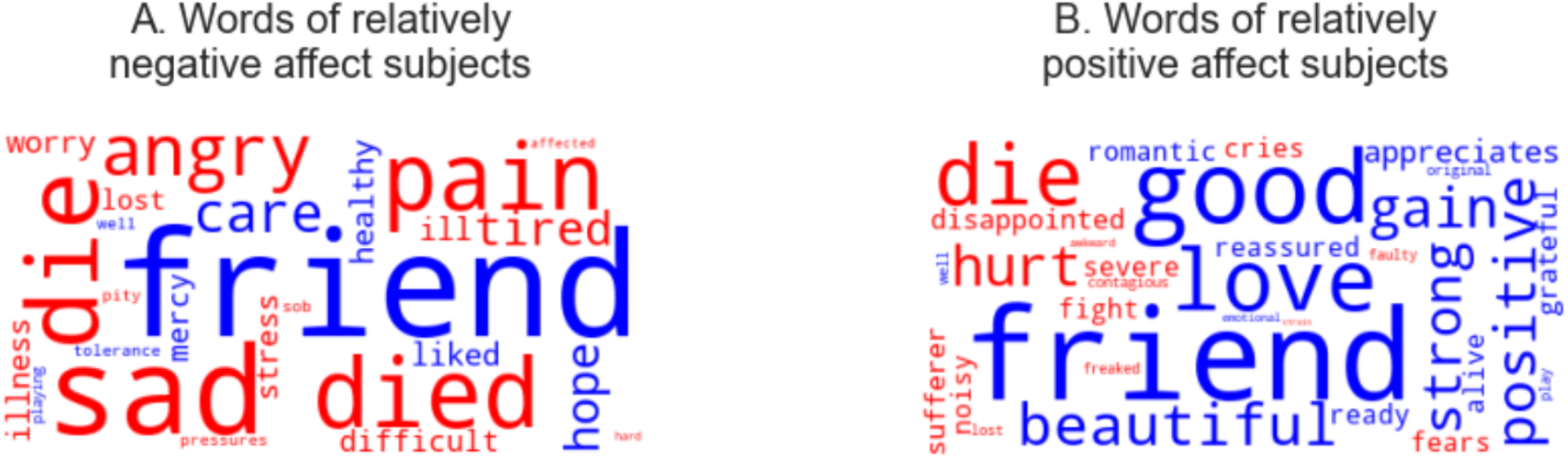
Word clouds showing words used by participants when recalling their social memories. A. Words used by subjects with the most negative social memory recall. B. Words used by subjects with the most positive social memory recall. Negative words are shown in red; positive words are shown in blue. Font size indicates strength of affect (highly negative or highly positive).

Importantly, there was no significant correlation (*r* = -.101, *p* = .536) or difference (t = -1.720, *p* = 0.093) in the affect scores between social (i.e., patient; Mean_social_ = -.148, SD = 1.894) and non-social (i.e., Khan Academy; Mean_nonsocial_ = .546, SD = 1.473) recall texts. These results suggest, respectively, that participants’ affect in their social memories is not conflated or redundant with their affect in their nonsocial memories, and that the affective content was “matched” for the social and non-social memories.

### On average, subjects’ functional connectivity similarity is high during baseline, encoding and consolidation phases

Before relating functional connectivity patterns to memory affect across subjects, we simply assessed the extent to which subjects exhibited similar functional connectivity profiles in default network regions (as well as limbic regions and across the whole brain) during baseline, encoding, and consolidation. As depicted in Fig. 3, this analysis requires first extracting, for each subject, the timecourse of neural activity from each default network region and correlating these timecourses between each pair of regions. Next, a vector is created for each subject, in which each vector cell is populated by the timecourse correlation of a default network ROI pair. Finally, all subjects’ vectors (i.e., network connectivity profiles) are correlated with one another. This analysis assesses the extent to which subjects’ functional connectivity profiles are similar to one another and helps ensure that across subjects, our measure of functional connectivity is reliable before examining its relationship to individual differences in memory content.

**Fig. 3.**
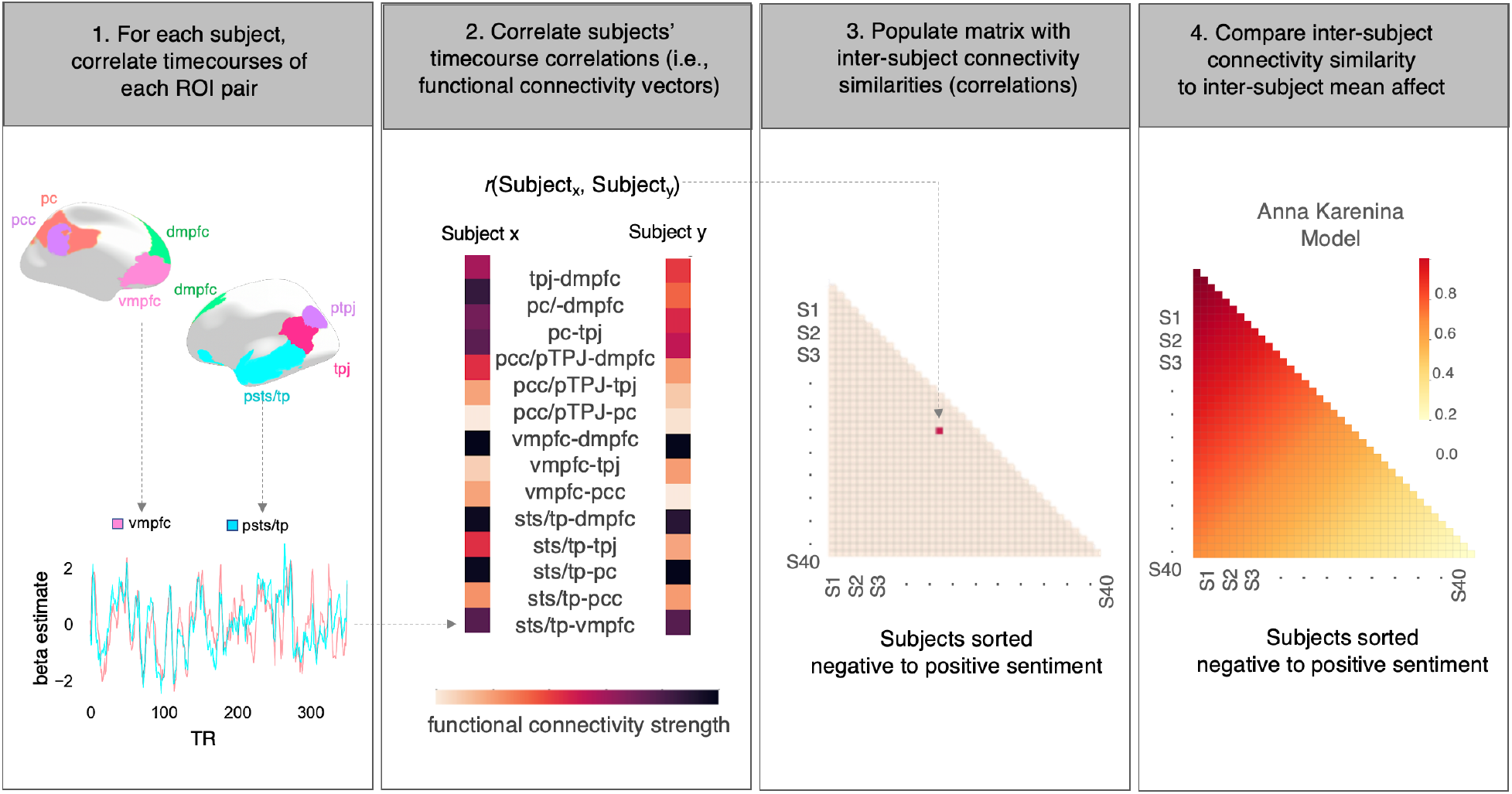
Data analytic approach involving four steps. First, for each subject and for each phase of the experiment, timeseries from the default network parcels were extracted and correlated with one another. Second, each pair of subjects’ default network functional connectivity vectors were correlated. Third, each subject pair’s connectivity similarity was populated into a subject-by-subject matrix. Fourth, the subject-by-subject connectivity similarity matrix was statistically compared to the theoretical Anna Karenina model. To examine the relative specificity of results to the default network, follow-up analyses were conducted with limbic region parcels, as well as all parcels throughout the brain.

Conceptually replicating prior work finding that neural synchrony is strong during naturalistic video watching^9,13,23^, functional connectivity similarity during social and non-social encoding was significant in the default network, between limbic regions (with and without NAc), as well as across the whole brain (Table 1). Interestingly, similarity in functional connectivity was also significant in these regions during baseline rest and social and non-social consolidation phases. This suggests that when collapsing across individual differences, subjects show similar functional connectivity profiles throughout the brain during each phase of the experiment.

**Table 1.** Across-subject similarity in functional connectivity was significant across phases (columns) and brain networks (rows).

|  | Baseline Rest |  | Social Encoding |  | Social Consolidation |  | Nonsocial Encoding |  | Nonsocial Consolidation |  |
| --- | --- | --- | --- | --- | --- | --- | --- | --- | --- | --- |
|  | *r* | *p* | *r* | *p* | *r* | *p* | *r* | *p* | *r* | *p* |
| Default Network | .559 | <.001 | .574 | <.001 | .523 | <.001 | .623 | <.001 | .475 | <.001 |
| Limbic Regions (without nucleus accumbens) | .974 | <.001 | .974 | <.001 | .933 | <.001 | .973 | <.001 | .958 | <.001 |
| Limbic Region (with nucleus accumbens) | .710 | <.001 | .710 | <.001 | .560 | <.001 | .705 | <.001 | .626 | <.001 |
| Whole Brain | .494 | <.001 | .565 | <.001 | .417 | <.001 | .553 | <.001 | .438 | <.001 |

Further, connectivity similarity was not significantly different during social consolidation compared to other phases (difference between social consolidation vs baseline: *r* = .047, *p* = .529; vs social encoding: *r* = .011, *p* = .498; vs non-social encoding: *r* < .001, *p* = .526; vs non-social consolidation: *r* = -.038, *p* = .472). Connectivity similarity was also not significantly different within the default network compared to limbic or whole brain networks (vs limbic_withoutNAc: *r* = .332, *p* = .488; vs limbic_withNAc: *r* = .032, *p* = .531; vs whole_brain: *r* = -.100, *p* = .517). Note that for these analyses, the r-value indicates dissimilarity, and thus significant r-values would indicate significantly distinct patterns (see Methods). These results suggest that any individual differences detected in subsequent analyses are not a function of, and are thus robust to, the underlying level of similarity across participants.

### During social consolidation, subjects with more negative social memories show similar default network functional connectivity, whereas subjects with more positive social memories show idiosyncratic default network functional connectivity: Inter-subject-Representational Similarity Analysis (IS-RSA)

Given our predictions about social consolidation processes during post encoding rest, we tested the Anna Karenina model on the social consolidation phase of the experiment. This step involves comparing the mean affect of each subject pair (‘inter-subject mean affect’) with their connectivity similarity (‘inter-subject connectivity similarity’). Fig. 3 provides a conceptual depiction of the IS-RSA analysis approach.

Note that the Anna Karenina model simultaneously tests the possibility that either 1) subjects with high negative recall show similar neural responding while subjects with high positive recall show dissimilar neural responding (which would be indicated by a positive correlation with the model in our formulation) or 2) subjects with high negative recall show dissimilar neural responding while subjects with high positive recall show similar neural responding (which would be indicated by a negative correlation with the model). The Anna Karenina model significantly predicted the affect in subjects’ social memories (*r* = .226, *p* = .029; Fig. 4A). This result indicates that subjects with highly negative social memories show similar default network functional connectivity profiles during consolidation, whereas subjects with highly positive memories show idiosyncratic default network functional connectivity patterns (i.e., their patterns are different from other positive subjects, as well as other negative subjects).

**Fig. 4.**
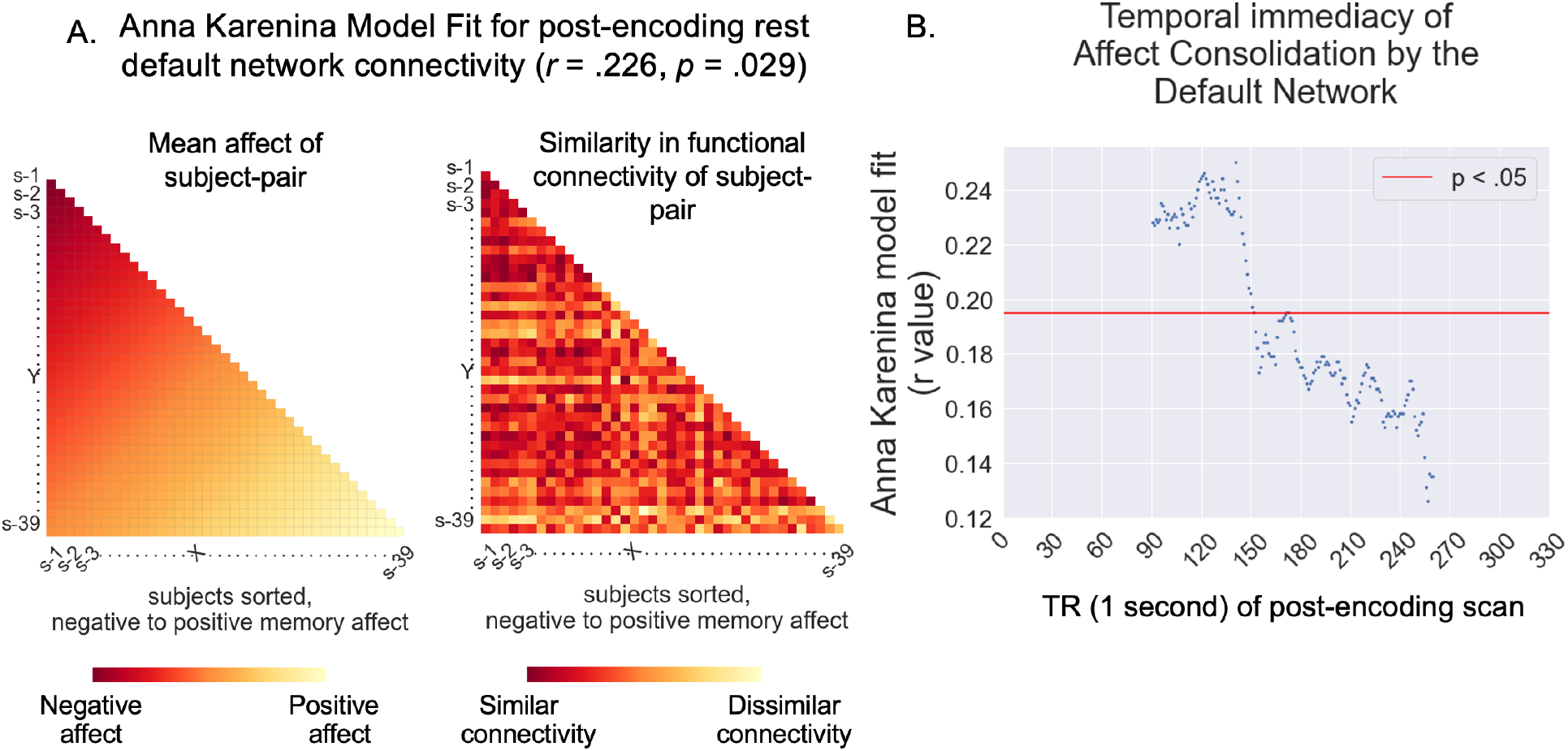
Panel A) shows that the Anna Karenina model significantly predicts the affect in subjects’ memories, with highly negative subjects showing similar default network functional connectivity profiles and highly positive subjects showing idiosyncratic default network functional connectivity profiles. Panel B) shows results from a sliding window analysis demonstrating that the Anna Karenina model is strongest and significant during earlier (vs. later) portions of the consolidation phase.

### IS-RSA results are only observed during the social memory consolidation phase

Previous research has identified that inter-subject correlation among default network regions stratifies according to individual differences during the *encoding* of an experience.^9,13,23^ It is thus possible to conceive that our consolidation results are driven by the remnants of functional connectivity patterns from the encoding phase. Moreover, given that subjects are resting in the scanner during memory consolidation, it is important to ensure that the consolidation IS-RSA results are not observed when subjects are resting in general (i.e., during the baseline resting state scan), as opposed to specifically during post-encoding rest. To rule out these alternatives, we repeated our analyses on default network functional connectivity profiles during encoding and baseline scans, finding no significant results during either phase (social encoding *r* = -.045, *p =*.646; baseline *r* = .013, *p* = .918). To further ensure that our findings are preferential to consolidating affect into *social* memories, relative to memories more generally, we next tested the Anna Karenina model on the *nonsocial* encoding and post-nonsocial-encoding consolidation phases and neither model was significant (nonsocial encoding *r* = -.018, *p* = .858; post-nonsocial-rest *r* = -.065, *p* = .511).

### IS-RSA results are not observed in limbic region or whole-brain analyses

Given the emotional nature of the videos, and the reliance of sentiment analysis on affective words, it is possible that other regions of the brain, such as those in the limbic system, would exhibit IS-RSA consolidation results in addition to the default network. To check for this, we ran our analyses on the consolidation phase in limbic regions associated with affective responding (*dACC, AI, and amygdala and separately, dACC, AI, amygdala, and nucleus accumbens*), as well as the entire brain. The Anna Karenina model was not significant for the limbic region analysis (*r*_*dACCAIamygdala*_ = -.123; *p* = .299; *r*_*dACCAIamygdalNAc*_ = -.150; *p* = .114) or whole brain analysis (*r* = .028; *p* = .779). We also examined whether subjects’ mean default network functional connectivity (i.e., the strength of their connectivity) correlated with affect scores, but this was not the case (*r* = - .142, *p*= .383). This further suggests that the default network effects we observed are driven by similarity in subjects’ functional connectivity profiles, as opposed to simply mean connectivity strength.

Finally, although affect scores were not significantly correlated with the number of correct facts recalled, to be certain that our Anna Karenina consolidation results are not related to the amount of information recalled, we ran an Anna Karenina model relating subjects’ number of correctly recalled facts with their default network functional connectivity profiles from the consolidation phase. This model was also non-significant (*r* = .005, *p* = .958), suggesting the observed affect results are not driven by memory accuracy more generally. Overall, our results converge to suggest that affect is consolidated into our social memories via default network functional connectivity during post-encoding rest, with subjects prone to negative memory consolidation showing similar default network profiles and subjects prone to positive memory consolidation showing idiosyncratic default network profiles.

### The role of specific regions-of-interest (ROIs)

Next, we were interested in probing which ROIs in the default network were driving our results. Specifically, we asked which ROIs’ exclusion from the network, individually or along with other ROIs, dissolved our results. To find this, we repeated our IS-RSA analyses with a default network region “subtraction method”, i.e., we considered a default network of size 5 where 1 of the 6 ROIs is dropped from the analyses, then a network of size 4 where every possible pair of ROIs is dropped from the analyses, then a network of size 3 where every possible triad of ROIs is dropped from the analyses. Our analyses yielded VMPFC as the only ROI that was necessary in all the permutations that showed significant effects. Among 5 node networks, the vmpfc-dmpfc-tpj.angular-precuneus-pcc *(r* = 0.247, *p* = 0.032), vmpfc-dmpfc-tpj.angular-precuneus-sts (*r* = 0.198, *p* = 0.016), and vmpfc-dmpfc-tpj.angular-pcc-sts *(r* = 0.206, *p* = 0.036) permutations showed significant effects; among 4 node networks, the vmpfc-dmpfc-tpj.angular-precuneus (*r* = 0.236, *p* = 0.008), vmpfc-dmpfc-pcc-sts (*r* = 0.228, *p* = 0.043), and vmpfc-tpj.angular-precuneus-sts (*r* = 0.191, *p* = 0.033) permutations showed significant effects; and among 3 node networks, the vmpfc-dmpfc-pcc (*r* = 0.209, *p* = 0.023) and vmpfc-tpj.angular-precuneus (*r* = 0.279, *p* = 0.016) showed significant effects. In other words, the default network requires the VMPFC to idiosyncratically consolidate positive affect and similarly consolidate negative affect into our social memories.

Additionally, given the established role of the hippocampus in memory formation^24^, we repeated our default network analyses while including the hippocampus in participants’ default network functional connectivity vectors. This model also significantly predicted subjects’ affect scores (*r* =.242, *p* =.019), suggesting that the default network works in concert with the hippocampus to consolidate individual differences in the affective content of social memories.

### IS-RSA consolidation results are strongest during early social memory consolidation

Finally, we investigated the temporal structure of our consolidation results. That is, is the relationship between default network functional connectivity and memory affect particularly salient during certain periods of memory consolidation? Considering the value of social memories in navigating everyday social relationships^25^ and their ability to ‘stick’ longer and resurface quicker than non-social memories^26^, and that default network regions engage “by default” during rest^53^, it has been speculated that social consolidation processes may occur relatively quickly by the default network.^27^ Thus, we next conducted our analyses separately on connectivity patterns during the first and second halves of the consolidation phase, finding significant effects only during the first half (first half of rest IS-RSA *r* = .232, *p* = .031; second half of rest IS-RSA *r* = .139 *p* = .196). Going one step further, we employed a sliding window approach, with each window still approximately half the duration of the consolidation phase (i.e., 3 minutes), to more precisely quantify when during rest the Anna Karenina model is most meaningful. To this end, the Anna Karenina model was tested across 3-minute time windows starting at t=0 and shifting by 1s each time. For example, the first 5 windows were 1s-180s, 2s-181s, 3s-182s, 4s-183s, and 5s-184s. We found a roughly linear reduction in our model fit, though it included a peak effect for the 50s-220s window before sharply dropping to non-significance by the 60s-230s mark (Fig. 4B). Analyses including the hippocampus showed the same profile of results. Overall, results suggest that individual differences in socioemotional memory consolidation occur relatively quickly during rest.

## Discussion

After interpersonal interactions, people often recall the event differently; some with optimism, some with despair; some are inspired, some deflated. Here we investigated how people who share the same social experience end up with negative versus positive memories. We discovered individual differences emerge during post-encoding rest in the brain’s default network. Specifically, individuals who end up with negative social memories gravitate towards similar default network functional connectivity patterns, while those with positive memories exhibit more idiosyncrasies. This is the first work to identify inter-subject (dis)similarities during consolidation at rest and adds to the growing area of research implicating the default network in interpersonal learning and memory. The findings also add novel support for theoretical accounts of asymmetric effects of affect on cognition^16^, including the broaden-and-build theory of affect^19,20^, which posits negative affect constricts, while positive affect broadens, the types of cognitive processes considered. For the first time, we identified i) post-encoding rest as a key moment and ii) the default network as a key brain system in which negative affect homogenizes, whereas positive affect diversifies cognitive processing.

The results are remarkably specific to the default network consolidating individual differences in the affective content of social memories during post-encoding rest. The Anna Karenina model, which tested the possibility that people who remember negative versus positive content may show different functional connectivity profiles, was only significant for default network regions (and not limbic regions or a whole brain analysis) and only during the social consolidation phase. The lack of IS-RSA results during baseline rest rules out the possibility that results are driven by persistent, trait-level differences in default network functional connectivity. The lack of IS-RSA results during social encoding further rules out the possibility that post social-encoding rest results are redundant with differences in perceptual processing *during* social experience. The lack of IS-RSA results during nonsocial encoding and consolidation phases further points to the specificity of the results to *social* learning. Moreover, participants’ *mean* default network functional connectivity during post-encoding rest was also unrelated to memory affect, further suggesting results pertain to inter-subject (dis)similarities in functional connectivity patterns.

Finally, the IS-RSA results are also highly specific to the affective content, rather than other, potentially conflated components of memory performance. For example, participants’ affect scores were not significantly correlated with the number of facts recalled from the videos and an IS-RSA model using the number of facts recalled was similarly non-significant. Additionally, we assessed the total number of words, the number of affective words, and the ‘affective intensity’ (i.e., the sum of absolute or unsigned affect scores) of participants’ recall, and none of these variables were associated with participants’ affect scores. These further rule out the possibility that our results are driven by alternative memory constructs. Collectively, the follow-up analyses converged to highlight post-social encoding rest as a key moment, and the default network as a key brain system, in driving individual differences in the affective content in social memories.

The results update and connect two previously siloed lines of research: one investigating memory consolidation during rest and the other examining how shared brain states relate to similarity in cognition and behavior. A great deal of research identifies post-encoding rest as a critical moment for memory.^4,28,29^ At the same time, a growing area of research investigates how neural similarity predicts meaningfully similar cognition and behavior.^9,30,31,32^ In fact, recent work in this area found that people who are closer to one another in a social network also show similar resting state functional connectivity profiles, particularly in the default network.^33^ Interestingly, “closeness” was defined in this study, in part, by people nominating whom they are likely to disclose information, and our paradigm requires that participants listen to others’ disclosures. Thus, the present findings are not only the first to show that inter-subject (dis)similarities extend to consolidation processes at rest; they further point to the possibility that default network consolidation processes at rest may play a key role in predicting whether and how social network members end up with similar “collective” memories in everyday social life.

It is noteworthy that while we did not observe a relationship between inter-subject differences *during* video watching and memory affect, previous research has identified encoding effects.

Priming and trait biases have been leveraged to evoke individual differences in default network while viewing ambiguous^12,32^ and polarizing^10,11^ stimuli, respectively. Highly emotional content has also been used to show that negative emotions correspond with greater inter-subject neural synchrony.^55^ However, the optimal trade-off between a stimulus’ ability to elicit individual differences versus synchrony is unknown^21^; this study, to our knowledge, is the first to use a stimuli which evokes synchrony during viewing while successfully eliciting individual differences only during the post-encoding phase. While future work is needed to determine if the inter-subject *encoding* effects observed in prior research carry over into post-encoding rest, we show that differences during post-encoding rest may nonetheless meaningfully explain differences in memory.

Hippocampus-default network relationships are often observed in cognitive neuroscience studies investigating memory. For example, slow-wave ripples in the hippocampus interact with default network activation during autobiographical memory recall.^34^ Likewise, we found that IS-RSA default network results during social memory consolidation persisted with the inclusion of the hippocampus. Our findings add novel insight to this body of research by identifying that hippocampal-default network coupling profiles during consolidation help explain individual differences in social memory. Future research using more invasive methods, such as intracranial electroencephalographic (iEEG) recordings^35^, may reveal the precise relationships between the hippocampus and default network in interpersonal memory consolidation.

Two sets of follow-up analyses revealed additional insight into the nature of the present results. First, temporal analyses revealed that our inter-subject results emerge during *early* post social encoding rest. The Anna Karenina model was significant during the first, but not second, half of the social consolidation phase and sliding window analyses further revealed that the effect occurs in the first ∼3.5 minutes of rest. The temporal immediacy may be driven by multiple factors. One possibility is that most consolidation effects occur during early rest, reflecting a type of recency effect in mind wandering. Alternatively, it is possible that interpersonal information, in particular, is temporally prioritized by the brain in the consolidation process. It has been argued that goal-relevant information may be “tagged” for prioritized memory consolidation at rest.^36^ Given that 1) humans have a strong, endogenous goal to feel connected to others^37^ and 2) default network regions engage “by default” during rest^53^, interpersonal information may be prioritized during post-encoding rest, with individual differences in affect codified quickly. Future work that manipulates when rest occurs after interpersonal interactions (e.g., immediately versus delayed) and 2) the duration of rest will clarify which of these competing possibilities best reflects how dis(similar) consolidation patterns generate individual differences in interpersonal memory.

The second set of follow-up analyses revealed VMPFC as a key player driving the default network results. We used a “subtraction” approach to the IS-RSA default network post-encoding rest analyses in which we considered how removing an ROI impacts results. This approach showed VMPFC as the only ROI that was necessary for the IS-RSA results. VMPFC is associated with generating affective interpretations, particularly positive ones.^38^ In addition to being a part of the default network, the VMPFC also works in concert with the ventral striatum to support positive affect.^39,40^ Moreover, recent work implicates VMPFC in supporting idiosyncratic affective responses^41^, particularly in response to naturalistic social stimuli.^42^ Our results therefore nicely complement and extend prior research on VMPFC, highlighting for the first time that this region plays a key role in asymmetrically tying negative and positive affect to social memories during rest.

More broadly, the results hint at a new way to think about the neurocognitive mechanisms supporting resilience and the “broaden and build” theory of positive emotion. In terms of resilience, extensive psychological research suggests that finding positive meaning in response to negative situations, such as a stressor bringing people together, promotes mental and physical health.^43^ While most of this research focuses on benefit finding among patients diagnosed with a disease, some work suggests that when caretakers find the benefits of a patient’s experience with a disease, the caretaker also experiences better well-being.^15^ Our paradigm mirrors this situation, as our participants listened to patients with cystic fibrosis share both negative and positive information about their diagnosis. This parallel, paired with the fact that prior work found VMPFC increases activity when participants are explicitly instructed to find positive meaning in response to negative stimuli^39^, points to the possibility that idiosyncratic VMPFC responding during consolidation may support the resilient strategy to see the good in the bad. Moreover, the broaden-and-build theory of positive emotion predicts that while negative emotions restrict thought patterns, positive emotions broaden the kinds of thoughts people entertain.^19,20^ Our results update both literatures, showing that benefit finding and the idiosyncratic thoughts that emerge with positive affect may occur spontaneously (i.e., without instruction) during post-encoding rest.

### Conclusion

What colors our social memories with positive versus negative hues? We find that individual differences in default network connectivity immediately after social experiences may explain why some of us recall them with optimism while others with despair. The results extend literature on asymmetric affect representation^19^ by showing that individuals with negative memories gravitate towards similar, while those with positive memories engage in unique thought patterns. At a broader level, these findings highlight the contribution of memory *consolidation* during rest – over and above encoding itself – in painting our social worlds with unique strokes.

## Methods

### Participants

Forty right-handed subjects (26 female; mean age = 29 years, SD = 11, 65% white; 23% Asian; 8% Hispanic) completed this study. Subjects either received $20 per hour of participation or were awarded course credit for completing the experiment. Subjects provided informed consent in accordance with the Dartmouth College institutional review board. The data used here has been reported on in prior work^54^, though notably all analyses reported here are orthogonal to those previously reported.

### Procedures

In the baseline and consolidation phases, participants saw a blank screen and were requested to rest but stay awake. Participants also completed an anatomical scan which was used for fMRI image processing.

### Sentiment analysis of memory recall

Each participant’s typed memory recall of the 4 social videos were combined into a recall paragraph per participant and the recall of the 4 nonsocial videos were combined into another recall paragraph per participant. The paragraphs were cleaned to exclude non-alphabets, common ‘stop-words’ (prepositions) as defined by python’s Natural Language ToolKit (NLTK)^44^, as well as neutral (non-affect words). Next, we performed sentiment analysis (VADER)^22^ on each word, such that negative words were affect scored between -1 and 0 (with more negative words scoring closer to -1), and positive words were affect scored between 0 and 1 (with more positive words scoring closer to 1). Finally, we summed the word affect scores across the entire paragraph to obtain one affect score per participant. We chose to use the sum instead of the average affect score because the variance and skew across summed affect scores yielded it to be the more appropriate metric to analyze (summed affect *var* = 3.68, *skew* = .135; mean affect *var* = -.004, *skew* = -.9). To rule out confounds, we ensured that the affect scores were not significantly correlated with the number of affect (*r* = .096, *p* = .554) or all words (*r* = .243, *p* = .131).

### fMRI Data Acquisition

Scanning was performed on a Siemens Prisma 3-T Trio. Functional images were acquired using an EPI gradient-echo sequence (2.5 × 2.5 × 2.5 mm voxels, repetition time = 1000 msec, echo time = 30 msec, 2.5-mm slice thickness, field of view = 24 cm, matrix = 96 × 96, flip angle = 59°, multiband acceleration factor = 4). A T2-weighted structural image was acquired coplanar with the functional images (0.9 × 0.9 × 0.9 mm voxels, repetition time = 2300 msec, echo time = 2.32 msec, 0.9-mm slice thickness, field of view = 24 cm, matrix = 256 × 256, flip angle = 8°).

### fMRI preprocessing

Functional and anatomical brain images were reoriented using SPM and skull-stripped using the Brain Extraction Tool in FSL. Data were preprocessed using FSL. Specifically, data underwent high-pass filtering (.009 Hz cutoff), motion correction, skull-stripping, spatial smoothing (6 mm radius), and registration to the anatomical image using Boundary-Based Registration. Nuisance variables, which included six standard motion parameters, their derivatives, as well as white matter and cerebrospinal fluid data, were regressed out using GLMs. Additionally, to correct for extreme motion, global (average brain) signal and motion scrubbing (volumes with framewise displacement > .2mm) artifacts were regressed out. All analyses are applied to the residual images from this nuisance-variable GLM.

### Neural time series extraction

We wanted to objectively define default network brain regions while also ensuring that the regions selected are functionally relevant to psychological constructs. For this reason, we used the k=50 whole brain parcellation that used k-means clustering to isolate meta-analytic coactivations^45^ from Neurosynth.^46^ This parcellation includes six parcels that comprise the default network: the dorsomedial prefrontal cortex (DMPFC), ventromedial prefrontal cortex (VMPFC), temporo-parietal junction (TPJ), precuneus (PC), posterior cingulate cortex functionally combined with posterior TPJ (pcc/pTPJ), and the superior temporal sulcus extending into temporal poles (STS/TP; default network regions depicted in Fig. 2). The whole brain parcellation also includes limbic regions traditionally associated with affective responding: dorsal anterior cingulate cortex (dACC), anterior insula (AI), amygdala, and nucleus accumbens (NAc). We therefore were able to examine functional connectivity (i.e., timecourse correlations) for each subject for 1) the default network, 2) limbic regions, and 3) across the whole brain (i.e., all 50 parcels). These and all subsequent analyses were performed on default network regions, as well as separately, brain regions in the limbic system as well as brain regions across the entire brain. We ran limbic region analyses two ways: first, with just the dACC, AI, and amygdala and second, with these regions as well as the nucleus accumbens (NAc), given its reliable role in positive affect. This two-pronged approach was taken because the dACC, AI, and amygdala are associated with both negative and positive affect^47^, whereas the NAc is more consistently associated with positive affect only.^48,49^

For each subject, the z-scored time series of neural activity in each parcel was extracted for each phase of scanning – baseline, social encoding, nonsocial encoding, social consolidation, and nonsocial consolidation. Following recommendations from prior work^50^, we not only excluded the few pre- and post-video fixation seconds, but also the first 10 seconds of the videos themselves, to prevent the onset of videos from inflating inter-subject similarity in connectivity. Similarly, we excluded the first 10 seconds of the baseline and consolidation phases each (i.e., the first 10 seconds of each rest period).

### Inter-subject similarity in Functional Connectivity

To calculate a participant’s default network functional connectivity, we computed the Pearson correlation of the neural time series between each default network parcel pair (separately for baseline, encoding, and consolidation), obtaining a 6*6 matrix of parcel-pair connectivity per participant. For inter-subject similarity analyses, default network functional connectivity vectors were created for each participant by vectorizing the lower triangle of a subject’s default network functional connectivity matrix (see Fig. 3). Specifically, following previous work’s methodology^33^, we measured the similarity in default network functional connectivity between participant pairs as *1 -Euclidean distance* of the pair’s z-scored functional connectivity vectors, obtaining an N*N inter-subject functional connectivity similarity matrix. We examined subjects’ functional connectivity patterns (rather than, for example, a given region’s timecourse) given prior work suggesting 1) consolidation during rest occurs via communication between brain regions^4^ and 2) when no stimuli are present as is the case during rest, between-participant similarity in functional connectivity profiles are more interpretable than between-participant similarity in regional timecourses.^33,51,52^

These steps were also taken in a series of follow-up analyses designed to assess the specificity of the results to default network regions. First, with the limbic regions dACC, AI, and amygdala, which created a 3*3 matrix of parcel-pair connectivity per subject, and then separately with nucleus accumbens (NAc), dACC, AI, and amygdala, which created a 4*4 matrix. Next, with the whole-brain parcellation which created a 50*50 matrix of parcel-pair connectivity per subject.

Finally, this step was also taken in a follow-up IS-RSA analysis that included the hippocampus (bilateral) with the default network regions. This created a 7*7 matrix of parcel-pair connectivity per subject. Given the hippocampus’s broad role in memory formation^24^, this analysis allowed us to further examine whether default network regions’ relationship with the hippocampus during post-encoding rest contributes to individual differences in affective memory content.

To test whether similarity in connectivity during social consolidation was significantly different from that during any of the other 4 phases (baseline, social encoding, non-social encoding, non-social consolidation), we conducted a non-parametric test in which we randomly sampled N=40 subjects, with repetition. We then generated the inter-subject connectivity similarity matrices for the same random subject sample for each of the 5 phases, thus ensuring that we permute at the subject (and not the subject-pair) level. The similarity of a subject with oneself, if present due to resampling, was discarded. Finally, we separately subtracted each phase’s inter-subject matrix from that of the social consolidation phase, to get 4 resultant difference matrices. The median value of each of these 4 difference matrices reflects the difference between social consolidation and each of the other 4 phases in the inter-subject connectivity similarity that they evoked. Repeating this 1000 times, we generated null distributions for the differences in inter-subject connectivity similarity between social consolidation and the other 4 phases. We then computed the p value as the number of times that the actual difference in inter-subject connectivity similarity between social consolidation and any other phase exceeds the null distribution of differences. Note that in this approach, the r-value indicates dissimilarity, and thus significant r-values would indicate significantly distinct patterns.

### IS-RSA Model creation and testing

We first converted subjects’ affect scores into ranks, such that negative subjects were ranked low and positive subjects were ranked high (range of ranks = 0-39 for N=40). Our Anna Karenina model stipulated a subject pair’s similarity in functional connectivity = 1-mean of the pair’s ranks, such that the higher the pair’s rank (indicating more positive recall), the lower their similarity in functional connectivity (indicating more idiosyncratic connectivity), and vice-versa. We thus obtained our 40*40 *connectivity similarity matrix model*. The Anna Karenina model is depicted in Fig. 3.

Finally, to test our hypothesis, we correlated our connectivity similarity matrix *model* with our connectivity similarity matrix *data* (we used spearman correlations to not assume linearity between the two matrices). To determine the statistical significance of a model’s fit, we needed to account for the non-independence in our data: specifically, each data point (matrix cell) represented a subject *pair*, and thus each *subject* was represented in multiple (N-1=39) data points. To this end, and consistent with prior work^6,9^, we conducted a non-parametric permutation test, wherein we randomly shuffled and reassigned subjects’ functional connectivity vectors 5000 times, each time correlating the resultant simulated matrix with our model matrix, thus generating a (null) distribution of IS-RSA (correlation) values. We then summed the number of times our simulated null correlation value exceeded our observed model-data correlation, yielding the probability that our results were generated by chance. Finally, we compared this probability against a significance threshold of alpha = .05 to discern statistical significance.

## Acknowledgments

This work was supported by NIMH R01MH125406 awarded to Dr. Meyer

## Code & Data Availability

Code & data are available at https://github.com/siyer7/default_network-socioaffective-memory-variability

## References

1. Everaert, J., Vrijsen, J. N., Martin-Willett, R., van de Kraats, L., & Joormann, J. (2022). A meta-analytic review of the relationship between explicit memory bias and depression: Depression features an explicit memory bias that persists beyond a depressive episode. Psychological Bulletin, 148(5-6), 435.

2. Durand, F., Isaac, C., & Januel, D. (2019). Emotional memory in post-traumatic stress disorder: A systematic PRISMA review of controlled studies. Frontiers in psychology, 10, 303.

3. Coles, M. E., & Heimberg, R. G. (2002). Memory biases in the anxiety disorders: Current status. Clinical psychology review, 22(4), 587–627.

4. Tambini, A., & Davachi, L. (2019). Awake reactivation of prior experiences consolidates memories and biases cognition. Trends in cognitive sciences, 23(10), 876–890.

5. Meyer, M. L., Davachi, L., Ochsner, K. N., & Lieberman, M. D. (2019). Evidence that default network connectivity during rest consolidates social information. Cerebral Cortex, 29(5), 1910–1920.

6. Chang, L. J., Gianaros, P. J., Manuck, S. B., Krishnan, A., & Wager, T. D. (2015). A sensitive and specific neural signature for picture-induced negative affect. PLoS biology, 13(6), e1002180.

7. Thornton, M. A., & Tamir, D. I. (2020). People represent mental states in terms of rationality, social impact, and valence: Validating the 3d Mind Model. Cortex, 125, 44–59.

8. Satpute, A. B., & Lindquist, K. A. (2019). The default mode network’s role in discrete emotion. Trends in cognitive sciences, 23(10), 851–864.

9. Finn, E. S., Corlett, P. R., Chen, G., Bandettini, P. A., & Constable, R. T. (2018). Trait paranoia shapes inter-subject synchrony in brain activity during an ambiguous social narrative. Nature communications, 9(1), 1–13.

10. Leong, Y. C., Chen, J., Willer, R., & Zaki, J. (2020). Conservative and liberal attitudes drive polarized neural responses to political content. Proceedings of the National Academy of Sciences, 117(44), 27731–27739.

11. van Baar, J. M., Halpern, D. J., & FeldmanHall, O. (2020). Intolerance to uncertainty modulates neural synchrony between political partisans. bioRxiv.

12. Simony, E., Honey, C. J., Chen, J., Lositsky, O., Yeshurun, Y., Wiesel, A., & Hasson, U. (2016). Dynamic reconfiguration of the default mode network during narrative comprehension. Nature communications, 7(1), 1–13.

13. Broom, T. W., Stahl, J. L., Ping, E. E., & Wagner, D. D. (2022). They saw a debate: Political polarization is associated with greater multivariate neural synchrony when viewing the opposing candidate speak. Journal of Cognitive Neuroscience, 35(1), 60–73.

14. Helgeson, V. S., Reynolds, K. A., & Tomich, P. L. (2006). A meta-analytic review of benefit finding and growth. Journal of consulting and clinical psychology, 74(5), 797.

15. Kim, Y., Schulz, R., & Carver, C. S. (2007). Benefit finding in the cancer caregiving experience. Psychosomatic medicine, 69(3), 283–291.

16. Fazio, R. H., Eiser, J. R., & Shook, N. J. (2004). Attitude formation through exploration: valence asymmetries. Journal of personality and social psychology, 87(3), 293.

17. Smallman, R., Becker, B., & Roese, N. J. (2014). Preferences for expressing preferences: People prefer finer evaluative distinctions for liked than disliked objects. Journal of Experimental Social Psychology, 52, 25–31.

18. Tugade, M. M., & Fredrickson, B. L. (2007). Regulation of positive emotions: Emotion regulation strategies that promote resilience. Journal of happiness studies, 8(3), 311–333.

19. Fredrickson, B. L. (1998). What good are positive emotions?. Review of general psychology, 2(3), 300–319.

20. Fredrickson, B. L. (2001). The role of positive emotions in positive psychology: The broaden-and-build theory of positive emotions. American psychologist, 56(3), 218.

21. Finn, E. S., Glerean, E., Khojandi, A. Y., Nielson, D., Molfese, P. J., Handwerker, D. A., & Bandettini, P. A. (2020). Idiosynchrony: From shared responses to individual differences during naturalistic neuroimaging. NeuroImage, 215, 116828.

22. Hutto, C., & Gilbert, E. (2014, May). Vader: A parsimonious rule-based model for sentiment analysis of social media text. In Proceedings of the international AAAI conference on web and social media (Vol. 8, No. 1, pp. 216–225).

23. Yeshurun, Y., Swanson, S., Simony, E., Chen, J., Lazaridi, C., Honey, C. J., & Hasson, U. (2017). Same story, different story: the neural representation of interpretive frameworks. Psychological science, 28(3), 307–319.

24. Bird, C. M., & Burgess, N. (2008). The hippocampus and memory: insights from spatial processing. Nature Reviews Neuroscience, 9(3), 182–194.

25. Murty, V. P., & Alison Adcock, R. (2017). Distinct medial temporal lobe network states as neural contexts for motivated memory formation. In The hippocampus from cells to systems (pp. 467–501). Springer, Cham.

26. Meyer, M. L. (2019). Social by default: characterizing the social functions of the resting brain. Current directions in psychological science, 28(4), 380–386.

27. Meyer, M. L. (2019). Social by default: characterizing the social functions of the resting brain. Current directions in psychological science, 28(4), 380–386.

28. Wamsley, E. J. (2019). Memory consolidation during waking rest. Trends in cognitive sciences, 23(3), 171–173.

29. McDonough, I. M., Letang, S. K., Erwin, H. B., & Kana, R. K. (2019). Evidence for maintained post-encoding memory consolidation across the adult lifespan revealed by network complexity. Entropy, 21(11), 1072.

30. Nummenmaa, L., Hari, R., Hietanen, J. K., & Glerean, E. (2018). Maps of subjective feelings. Proceedings of the National Academy of Sciences, 115(37), 9198–9203.

31. Stephens, G. J., Silbert, L. J., & Hasson, U. (2010). Speaker–listener neural coupling underlies successful communication. Proceedings of the National Academy of Sciences, 107(32), 14425–14430.

32. Sievers, B. R., Welker, C. Hasson, U., Kleinbaum, A. M. & Wheatley, T. (invited revision). How consensus-building conversation changes our minds and aligns our brains.

33. Hyon, R., Youm, Y., Kim, J., Chey, J., Kwak, S., & Parkinson, C. (2020). Similarity in functional brain connectivity at rest predicts interpersonal closeness in the social network of an entire village. Proceedings of the National Academy of Sciences, 117(52), 33149–33160.

34. Norman, Y., Raccah, O., Liu, S., Parvizi, J., & Malach, R. (2021). Hippocampal ripples and their coordinated dialogue with the default mode network during recent and remote recollection. Neuron, 109(17), 2767–2780.

35. Parvizi, J., & Kastner, S. (2018). Promises and limitations of human intracranial electroencephalography. Nature neuroscience, 21(4), 474–483.

36. Cowan, E. T., Schapiro, A. C., Dunsmoor, J. E., & Murty, V. P. (2021). Memory consolidation as an adaptive process. Psychonomic Bulletin & Review, 28(6), 1796–1810.

37. Baumeister, R. F., & Leary, M. R. (1995). The need to belong: Desire for interpersonal attachments as a fundamental human motivation. Psychological Bulletin, 117(3), 497–529.

38. Roy, M., Shohamy, D., & Wager, T. D. (2012). Ventromedial prefrontal-subcortical systems and the generation of affective meaning. Trends in cognitive sciences, 16(3), 147–156.

39. Doré, B. P., Boccagno, C., Burr, D., Hubbard, A., Long, K., Weber, J., … & Ochsner, K. N. (2017). Finding positive meaning in negative experiences engages ventral striatal and ventromedial prefrontal regions associated with reward valuation. Journal of cognitive neuroscience, 29(2), 235–244.

40. Lin, W. J., Horner, A. J., & Burgess, N. (2016). Ventromedial prefrontal cortex, adding value to autobiographical memories. Scientific reports, 6(1), 1–10.

41. Lieberman, M. D., Straccia, M. A., Meyer, M. L., Du, M., & Tan, K. M. (2019). Social, self,(situational), and affective processes in medial prefrontal cortex (MPFC): Causal, multivariate, and reverse inference evidence. Neuroscience & Biobehavioral Reviews, 99, 311–328.

42. Chang, L. J., Jolly, E., Cheong, J. H., Rapuano, K. M., Greenstein, N., Chen, P. H. A., & Manning, J. R. (2021). Endogenous variation in ventromedial prefrontal cortex state dynamics during naturalistic viewing reflects affective experience. Science Advances, 7(17), eabf7129.

43. Helgeson, V. S., Reynolds, K. A., & Tomich, P. L. (2006). A meta-analytic review of benefit finding and growth. Journal of consulting and clinical psychology, 74(5), 797.

44. Bird, S. 2006, July). NLTK: the natural language toolkit. In Proceedings of the COLING/ACL 2006 Interactive Presentation Sessions (pp. 69–72).

45. de la Vega, A., Chang, L. J., Banich, M. T., Wager, T. D., & Yarkoni, T. (2016). Large-scale meta-analysis of human medial frontal cortex reveals tripartite functional organization. Journal of Neuroscience, 36(24), 6553–6562.

46. Yarkoni, T., Poldrack, R. A., Nichols, T. E., Van Essen, D. C., & Wager, T. D. (2011). Large-scale automated synthesis of human functional neuroimaging data. Nature methods, 8(8), 665–670.

47. Cunningham, W. A., & Brosch, T. (2012). Motivational salience: Amygdala tuning from traits, needs, values, and goals. Current Directions in Psychological Science, 21(1), 54–59.

48. Knutson, B., Adams, C. M., Fong, G. W., & Hommer, D. (2001). Anticipation of increasing monetary reward selectively recruits nucleus accumbens. Journal of Neuroscience, 21(16), RC159–RC159.

49. Koch, M., Schmid, A., & Schnitzler, H. U. (1996). Pleasure-attenuation of startle is disrupted by lesions of the nucleus accumbens. Neuroreport, 7(8), 1442–1446.

50. Nastase, S. A., Gazzola, V., Hasson, U., & Keysers, C. (2019). Measuring shared responses across subjects using inter-subject correlation. Social Cognitive and Affective Neuroscience, 14(6), 667–685.

51. Mars, R. B., Verhagen, L., Gladwin, T. E., Neubert, F. X., Sallet, J., & Rushworth, M. F. (2016). Comparing brains by matching connectivity profiles. Neuroscience & Biobehavioral Reviews, 60, 90–97.

52. Liu, W., Kohn, N., & Fernández, G. (2019). Inter-subject similarity of personality is associated with inter-subject similarity of brain connectivity patterns. Neuroimage, 186, 56–69.

53. Raichle, M. E., MacLeod, A. M., Snyder, A. Z., Powers, W. J., Gusnard, D. A., & Shulman, G. L. (2001). A default mode of brain function. Proceedings of the national academy of sciences, 98(2), 676–682.

54. Collier, E., & Meyer, M. L. (2020). Memory of others’ disclosures is consolidated during rest and associated with providing support: Neural and linguistic evidence. Journal of Cognitive Neuroscience, 32(9), 1672–1687.

55. Nummenmaa, L., Glerean, E., Viinikainen, M., Jääskeläinen, I. P., Hari, R., & Sams, M. (2012). Emotions promote social interaction by synchronizing brain activity across individuals. Proceedings of the National Academy of Sciences, 109(24), 9599–9604.

